# Development of a microfluidic cell culture and monitoring system for intracellular signaling studies

**DOI:** 10.1101/453100

**Authors:** Tomoki Ohkubo, Haruyuki Kinoshita, Toshiro Maekawa, Katsuyuki Kunida, Hiroshi Kimura, Shinya Kuroda, Teruo Fujii

## Abstract

We describe a microfluidic cell culture and monitoring system that temporally controls molecule concentrations around cells cultured in a small space. The simple system consists of three syringe pumps and a microfluidic device with two inlet ports and two outlet ports. Each syringe pump discharges or draws culture medium, solutions containing signal molecules, or cell suspensions through a port in a programmed flow rate sequence. Signal molecule solutions of differing concentration are merged in a microchannel, mixed immediately, and transported into the cell culture chamber. Regulating the flow rate ratio of syringe pumps over time enables dynamic control of the concentration of signal molecules in the cell culture chamber. The system provides various time-dependent waveforms of concentration over cultured cells, including pulse, rectangular, and triangular. The practical performance of the system for concentration control was evaluated using fluorescent dye imaging. The system was also used with CHO-K1 cells to measure intracellular Ca^2+^ concentrations, which vary with extracellular ATP levels. When a rectangular pulse of ATP was applied to the cells, Ca^2+^ levels increased quickly. By contrast, several Ca^2+^ peaks were observed in response to stepwise increases in ATP concentration. Single-cell Ca^2+^ responses to ATP pulse stimulation were analyzed by quantitative fluorescence imaging. Hierarchical clustering and quantitative analysis of single-cell data revealed the diversity of Ca^2+^ responses to ATP pulse stimulation. These results demonstrate that the microfluidic cell culture system is useful for studying a variety of cellular responses, including cell signaling.

## Introduction

Processes in the cellular microenvironment are highly time-dependent due to the presence of soluble factors such as cytokines, hormones, growth factors, and neurotransmitters, the concentrations of which change dynamically. Recent studies have reported that some biological information is encoded in the temporal profiles of soluble factor concentrations *in vivo* for cell-cell communication^1–5^. For example, hepatocellular metabolism is regulated by temporal changes in extracellular insulin concentrations^6,7^. Elucidating the relationships between dynamic extracellular stimuli and resulting cellular responses is a key to understanding the dynamics of intracellular signaling mechanisms.

*In vitro* cell culture is a key technology used to monitor cellular responses to dynamic changes in the chemical environment, as it enables precise control of environmental conditions. In order to deepen understanding of cellular dynamics, an *in vitro* technique to accurately produce variations in temporal patterns of soluble factor concentrations is needed. Unfortunately, conventional cell culture methods based on pipetting and traditional plastic culture ware are not well-suited to producing temporal changes in concentration with a high degree of accuracy. Also, controlling environmental conditions while imaging the responses of live cells is difficult using conventional cell culture methods. Therefore, this study focused on the development of a cell culture system that enables both automatic perfusion and flexible stimulus control to facilitate quantitative investigations of the mechanisms of encoding/decoding systems (which we denote “temporal coding”) via signaling pathways.

Microfluidics, a set of technologies that enable fabrication of micro-scale structures and handling of microscopic fluid flows, holds tremendous promise for advancing biological research. The unique characteristics of microfluidics, such as fine geometric patterns, laminar liquid flow, and mixing purely based on diffusion, provides a diverse range of fluidic control strategies. A number of methods have recently been developed that enable modulation of the extracellular environment via dynamic control of the concentration of signal molecule^8–16^.

In order to generate temporal variation of signal molecule concentration around cells, a typical microfluidic system delivers a solution containing the molecule of interest and a diluent into a mixing channel, where they are mixed to homogeneity via molecular diffusion prior to delivery to the cells. To produce patterns of temporal variation in concentration, the mixing ratio of the two solutions is varied with time while maintaining a constant total flow rate. Several reports have described cell culture devices that enable the generation of multiple types of waveforms of glucose concentration, such as square, triangular, and sinusoidal waves^17–20^. In these devices, the molecule of interest and a diluent are pumped into a mixing channel by sequentially switching a series of on-chip diaphragm valves. The flow rate ratio of the two solutions is accurately controlled by careful and precise operation of the valves. More elaborate techniques have also been developed to produce temporal waveforms. For example, pulse code modulation was used to produce automated temporal changes in the concentrations of stimulants^13,21–23^. In their technique, off-chip pumps deliver the stimulant and a diluent to a microfluidic device. A series of on-chip valves pulsate the streams of the two solutions at given durations and a specified number of duty cycles in a mixing channel, which accelerates longitudinal diffusion. The concentration of stimulant can be regulated by changing the duration and number of duty cycles.

The main disadvantage of the above-mentioned pumping methods based on sequential switching of on-chip valves or pulse code modulation is their design complexity. Numerous tubes must be connected between the microfluidic device and pressure source to drive the on-chip valves and pumps, which increases the risk of dropout and tube clogging. Moreover, these systems are often expensive. For cell culture, almost all devices should be single use. We have previously reported a microfluidic device composed of a fluid driving unit, channels, and reservoirs^24^, however, these existing devices employ a complex channel design or multilayer structure comprised of a thin membrane of silicone elastomer and etched glass; such devices are too expensive to be disposed of after use. Thus, there is great need for a much simpler and more robust method for accurately generating temporal variation of signal molecule concentration and delivering them to cultured cells.

This work aimed to develop a microfluidic cell culture system employing a straightforward method to produce patterns of signal molecule concentration that vary over time. The system described is composed of three main components: 1) a disposable microfluidic device, 2) high-precision syringe pumps for concentration control and cell/reagent introduction, and 3) a universal fluidic interface linking the syringe pumps to the device. The developed system is capable of producing different types of concentration waveforms (e.g., step, triangular, and rectangular) and precisely regulating slope of those waveforms. For example, a gradual slope is necessary for a triangular wave gradient, whereas step or rectangular waves require steep rises and falls. To meet these requirements, the microchannel geometry was designed based on hydrodynamics. A microfluidic device was then manufactured using a standard microfabrication process called softlithography. A prototype of the microfluidic cell culture system was constructed by assembling the developed microfluidic device, high-precision syringe pumps, and other components. The developed system enables visualization of fluid flow in the microfluidic device by incorporating a fluorescent dye into the medium as a visible substitute for molecules of interest, and the concentration can be measured in the cell culture chamber.

The developed cell culture system was applied to the measurement of intracellular Ca^2+^ concentrations in Chinese hamster ovary (CHO-K1) cells, which are known to exhibit a rapid response to changes in the concentration of adenosine triphosphate (ATP). The Ca^2+^ responses of single cells to an ATP pulse stimulus were quantitatively analyzed by fluorescence imaging. The diversity of Ca^2+^ responses to ATP pulse stimulation were characterized by hierarchical clustering and quantitative analysis of the single-cell data. The ability to characterize responses of CHO-K1 cells to extracellular ATP concentration demonstrates that the developed system is useful for the study of cellular signaling mechanisms.

## Experimental

### Design and fabrication

The developed microfluidic cell culture system is illustrated in Figure 1. The system consists of a simple PDMS/glass microfluidic device and three syringe pumps (MFS-SP1, Microfluidic Systems Works Inc.), each of which is equipped with a glass syringe (gastight syringe 1001RN 1.0 mL, Hamilton Company) (Fig. 1a). The microfluidic device is mounted on the stage of an inverted epifluorescence microscope and fixed by a pair of microfluidic interface modules, which enables quick and easy tube connection and minimizes the dead volumes between the device and the connecting tubes to reduce hydrodynamic damping effects. One of the syringe pumps (Pump A) discharges the signal molecule solution at a constant flow rate, and another syringe pump (Pump B) discharges the diluent solution (signal molecule (−) in Fig.1a). The two solutions are pumped into the inlet ports of the Y-shaped microchannel and mixed in the mixing channel, which has a high-aspect-ratio rectangular cross section to enhance mixing by molecular diffusion at the liquid-liquid interface. A third syringe pump (Pump C) is used to withdraw liquid from the cell culture chamber. This suction pump is used to introduce cell suspension or fluorescent labeling reagents into the cell culture chamber from a pipette tip inserted into the outlet port of the interface module as a reservoir as depicted in Figure 2. In addition, the slight suction flow produced by Pump C helps stabilize the flow field in the device. The flow rates of Pumps A and B are controlled using a personal computer and microcontroller (Arduino MEGA 2560, Arduino LCC) with a programmed sequence. Pump C is operated manually on demand.

**Fig. 1.**
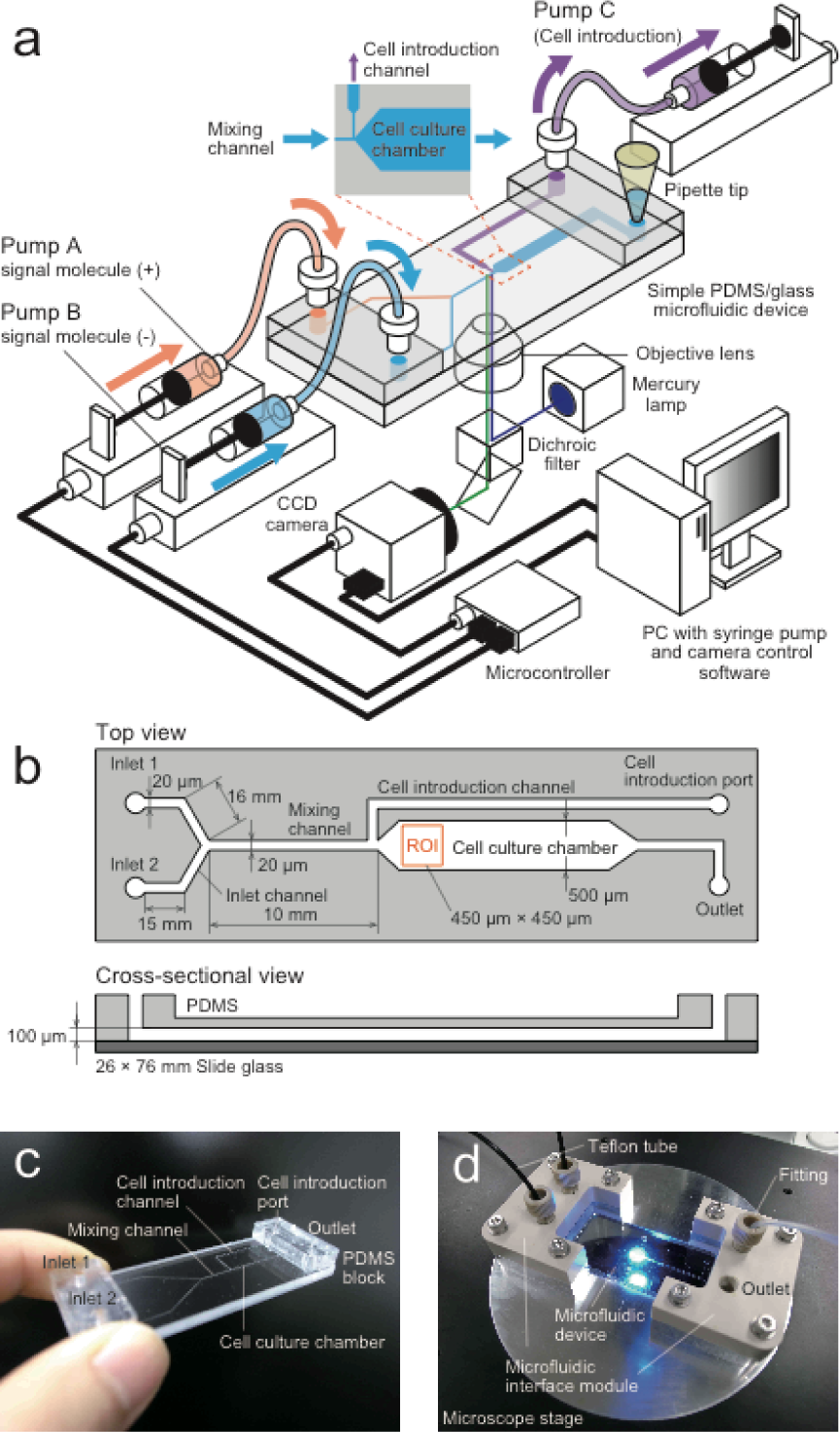
Microfluidic cell culture system (drawings are not to scale). (a) Schematic illustration of the system including three syringe pumps, a microfluidic device, an imaging unit, and a control system. (b) Top and side views of the microchannel design. Two solutions are merged and mixed in the mixing channel prior to entering the cell culture chamber. Cell suspensions and reagents are introduced from the cell introduction channel into the cell culture chamber. (c) Photograph of the microfluidic device. A pair of PDMS blocks with two holes is bonded at the inlet and outlet ports of the device for tubing. (d) Photograph of the microfluidic interface modules. The modules are used not only to connect tubes but also to hold the microfluidic device on the stage of an inverted microscope.

**Fig. 2.**
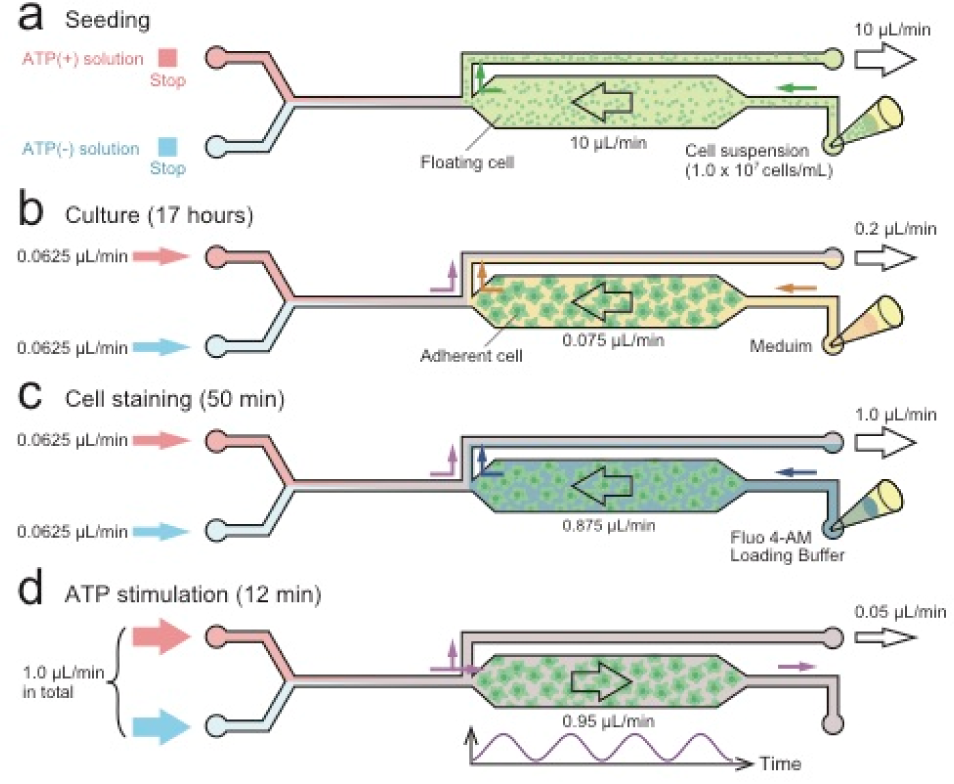
Procedure for ATP stimulation. Seeding, culturing, cell staining, and ATP stimulation are performed by operating three syringe pumps. (a) Cell seeding. Cell suspension was introduced from the reservoir pipette tip into the cell culture chamber at 10 µL/min by drawing it with Pump C. (b) Cell culture. The medium was supplied from the reservoir pipette tip into the cell chamber at 0.075 µL/min by operating Pump C. Pumps A and B were also feeding solutions at 0.0625 µL/min to stabilize the flow field. (c) Cell staining. Fluo 4-AM loading buffer was supplied from the reservoir pipette tip into the cell culture chamber at 1.0 µL/min in the same way as the cell culture. Cells were exposed to the Fluo 4-AM loading buffer for 50 min. (d) ATP stimulation. Pumps A and B were started in sequence to generate a concentration waveform. To prevent backflow from the cell introduction channel into the cell culture chamber, Pump C continued drawing at a 0.05 µL/min.

The dimensions of the microchannels are designed to theoretically improve the hydrodynamic response, prevent backflow, and reduce hydrodynamic shear stress on cells. Hydrodynamic shear stress and pressure drops were estimated based on the theoretical equation describing laminar flow in a square channel. The microfluidic device is illustrated in Figure 1b. The two incoming fluid flows are merged at the Y-junction, mixed in a mixing channel, and transported into the cell culture chamber. The size of the cell culture chamber is 500 µm × 100 µm × 20 mm (W × H × L). In all subsequent experiments described below, only cells located in a 450-µm square region of interest (ROI) in the cell culture chamber were evaluated (Fig.1b). The total flow rate of Pumps A and B,*Q*_total_, was kept constant at 1.0 µL/min. By contrast, the suction flow rate of Pump C was set at 0.05 µL/min. Under these flow rate conditions, it was estimated that the liquid inside the ROI would be replaced within a few seconds. The mean flow velocity in the cell culture chamber was 316 µm/s, resulting in wall shear stress of 0.15 dynes/cm^2^ on the bottom surface of the cell culture chamber. This means that the cells in the cell culture chamber are exposed to a shear stress of 0.15 dynes/cm^2^ during experiments, which is sufficiently small to have a negative effect on cell adhesion, growth, differentiation, or signal transduction^25^.

The mixing channel is a key component, as it forms a homogeneous distribution of the signal molecule in the cell culture chamber. The dimensions (W × H × L) of the mixing channel are 20 µm × 100 µm × 10 mm. In order to complete mixing of two solutions in a microchannel, the channel width should be shorter than twice the average molecular diffusion length. The time required for complete mixing is expressed by *W*^2^/4σ using the diffusion coefficient of a solute molecule, σ. By comparison, *Whl/Q_total_* represents the time required for the molecules to pass through the mixing channel with the fluid flow. Therefore, for perfect mixing, the dimensions of the mixing channel must satisfy the following condition:

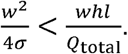

According to this theoretical analysis, the above-described mixing channel can homogeneously mix molecules with a diffusion coefficient of _≤_83.3 µm^2^/s.

Another significant hydrodynamic function of the developed microfluidic device is to prevent backflow at the Y-shaped junction from the mixing channel. For this purpose, the two merging channels at the junction were designed to have higher hydraulic resistance than the mixing channel by reducing their width. Both merging channels have the same cross section (W20 µm × H100 µm) and length (31 mm). The hydraulic resistance of each merging channel was estimated as 7.0 kPa, which is high enough to prevent backflow from the mixing channel, given that its hydraulic resistance is 2.2 kPa.

The microfluidic device was fabricated by softlithography using a negative photoresist and replica molding (Fig. 1c). The microchannel pattern was drawn using AUTOCAD (Autodesk Inc.) and printed on a Cr glass blank mask (Clean Surface Technology Co., Ltd.) at high resolution (minimum line and space < 1 µm) using a laser maskless lithography system (DWL40, Heidelberg Instruments Mikrotechnik GmbH). To develop high-aspect-ratio structures, KMPR 1035 photoresist (MicroChem Corp.) was directly spin-coated on the Cr surface of the printed mask to 100-µm thickness. The photoresist layer was exposed to UV light from the glass surface of the mask. After development of the KMPR1035 mold, PDMS (polydimethylsiloxane; Silpot184, Dow Corning Toray Co., Ltd.) was cast to a thickness of 2 mm, cured at 75°C for 1.5 h, and then permanently bonded to a slide glass (Matsunami Glass Ind., Ltd.) by irradiation of oxygen plasma using a reactive ion etching system (RIE 10NR, Samco, Inc.). A pair of PDMS blocks with two access holes for connection to the microfluidic interface modules were made by PDMS casting and bonded to the inlet and outlet ports of the fabricated microfluidic device after oxygen plasma treatment. The device was heated at 75°C for 30 min to ensure the strength of bonding. A pair of universal fluidic interfaces was set on the inlet and outlet ports of the microfluidic device to link the syringe pumps to the device (Fig. 1d).

### Concentration regulation

The performance of the developed system in terms of concentration regulation was demonstrated and evaluated through visualization and measurement of a fluorescent dye (FITC I dextran, 70 kDa, Research Organics, Inc.) in distilled water as a visible substitute for a signal molecule. The concentration of fluorescent dye solution in Pump A was 5 mg/mL. The total Pump A (dye solution) and Pump B (water) flow rate was set at 1.0 µL/min, and the suction flow rate of Pump C was set at 0.05 µL/min. The flow rate ratio between Pump A and B was regulated to achieve the desired mixing ratio in each experiment.

The developed microfluidic system was mounted onto the stage of an inverted epifluorescence microscope (IX71, Olympus Corp.) equipped with an objective lens (UPLFLN 4X, NA 0.13, Olympus Corp.), an 8-bit monochromatic CCD camera (DMK31BU03, The Imaging Source Europe, GmbH), and a fluorescent filter cube (U-NIB3, Olympus Corp.). The frame rate and exposure time for imaging were set at 3.76 fps and 1/94 s, respectively. Sequential 100-frame, 8-bit grayscale images were captured by this system and obtained 100 images were averaged to eliminate pixel flickering. The resulting averaged image was analyzed using ImageJ software (National Institutes of Health) to extract the fluorescence intensity of each pixel in the ROI.

Spatial uniformity of dye distribution in the cell chamber was evaluated by quantifying spatial variation of fluorescence intensity of the dye in the ROI. The coefficient of variation (CV) of grayscale intensity in the ROI was used as an index of spatial dispersion of concentration^26–30^ in this study. The CV was defined by the following equation:

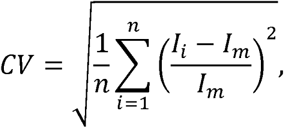

where *I_i_* represents the grayscale intensity at a given pixel in the ROI, *I_m_* represents the averaged intensity over the ROI, and *n* represents the total number of pixels in the ROI. Using this definition, CV = 0 means that the dye molecules are uniformly distributed in the cell chamber. By contrast, CV = 1 indicates that the dye solution and the water are streaming separately in the flow field of the cell chamber without mixing. The CVs in the ROI for different flow rate ratios were measured using the developed system. The CVs of standard solutions, which were homogenously premixed, were also measured for each mixing ratio as references. Standard solutions of the dye (FITC I dextran, 70 kDa) were prepared at 17 different concentration over the range 0 to 5.0 mg/mL and introduced sequentially into the cell culture chamber. The minimum detection limit CV was estimated by measured CVs in the developed imaging system.

A calibration curve showing the relationship between dye concentration and detected fluorescence intensity is essential for measuring the dye concentration quantitatively. To construct the calibration curve, the standard solutions of dye (FITC I dextran, 70 kDa) prepared for evaluating mixing uniformity were used. One of the standard dye solutions was pumped into the cell chamber from a reservoir pipette tip at a constant flow rate of 1.0 µL/min and imaged under the above-mentioned conditions. The mean fluorescence intensity in 36 images of the ROI was then calculated. This analysis was repeated three times for each standard solution to ensure reproducibility. Finally, the values of fluorescence intensity and dye concentration were plotted and fitted to an exponential function to construct the calibration curve.

Two primary types of concentration waveforms (rectangular and triangular waves) were produced in the cell culture chamber to demonstrate that the developed system can regulate the molecule concentration temporally in a programmed sequence. The dye concentration in the cell culture chamber was fluctuated sharply up and down in the rectangular wave by repeating on-off operation of Pumps A and B at constant time intervals. In this experiment, the flow rates of both pumps were switched between 0 and 1.0 µL/min at 90‐ or 180-s intervals. Gradual increase and decrease of the concentration as in a triangular wave were accomplished by changing the flow rate ratio between Pumps A and B. To produce an upslope concentration profile, the flow rate of Pump A was increased in steps of 1/16 µL/min from 0 to 1.0 µL/min, whereas the flow rate of Pump B was decreased oppositely in the same steps from 1.0 to 0 µL/min. A downslope profile was also formed by varying the acceleration/deceleration of each pump. In this experiment, the upslope and downslope operations were repeated alternately several times at 90‐ or 180-s intervals to demonstrate a triangular wave profile. The mean fluorescence intensity in the ROI was extracted from each fluorescence image, converted to a concentration value using the calibration curve, and then plotted.

### Cell stimulation and analysis

In order to verify the performance of the developed cell culture system for studying cellular responses to dynamically changing chemical environments, changes in Ca^2+^ concentrations in CHO-K1 cells were examined. CHO-K1 cells show time-dependent responses to the concentration of extracellular ATP. Fluo 4-AM (Calcium Kit Fluo 4-AM, Dojindo Molecular Technologies, Inc.), a green fluorescent Ca^2+^ probe, was used to visualize changes in intracellular Ca^2+^ concentrations. This probe emits green fluorescence with a peak wavelength of 518 nm upon exposure to Ca^2+^ ions. Extracellular ATP served as a messenger to induce increases in intracellular Ca^2+^ concentrations in this experiment. ATP was premixed with the Recording Medium included in the Calcium Kit Fluo 4-AM at a concentration of 1,000 nM. A red fluorescent dye (rhodamine B isothiocyanate-dextran, average molecular weight 10,000, Sigma-Aldrich Corp.) was also added to the ATP solution to visualize and quantify the ATP concentration via the emission of red fluorescence with a peak wavelength of 625 nm. The ATP concentration was indirectly estimated by visualizing and measuring the fluorescence intensity of the red dye, as it has almost the same diffusion coefficient as ATP molecules.

During the experiments, the cells in the microfluidic device were incubated in a CCM-1.3XYZ/CO_2_ real-time cultured cell monitoring system (Astec Co., Ltd.) to control the culture environment and permit live imaging of the cells. The system provides favorable environments for cells, such as 5% CO_2_, ambient temperature of 37°C, and sufficient humidity. Another key feature of this system is automated time-lapse fluorescence microscopy imaging. An epifluorescence microscope unit is installed in the system, which enables imaging of two different fluorescent markers (Fluo 4 and rhodamine B) alternately at 10-s intervals in the cell culture environment.

Figure 2 illustrates the procedures of the ATP stimulation experiment. The surface of the cell culture chamber was coated with collagen in advance by injecting a collagen solution (Cellmatrix Type I-C, Nitta Gelatin, Inc.) into the device, incubating it at 37°C for 30 min, and then rinsing with medium. CHO-K1 cells were obtained from Cell Bank (RIKEN BioResource Center) and cultured in Dulbecco’s Modified Eagle’s medium (low glucose, pyruvate; Life Technologies Corp.) supplemented with 10% fetal bovine serum (Sigma-Aldrich Corp.) in a tissue culture dish at 37°C in a humidified 5% CO_2_ incubator. The cells were then dissociated from the dish with 0.25% Gibco trypsin/EDTA (Thermo Fisher Scientific, Inc.) and collected in a centrifuge tube. The collected cells were centrifuged and resuspended at a concentration of 10^7^ cells/mL and introduced from the reservoir pipette tip into the cell culture chamber of the device by operating Pump C at 10 µL/min (Fig. 2a). After 30 min to allow for cell adhesion, perfusion culture was started by replacing the reservoir pipette tip with a tip filled with fresh medium and activated Pump C to draw the medium into the cell culture chamber at 0.2 µL/min. Simultaneously, Pumps A and B were started to feed their solutions at 0.0625 µL/min respectively to stabilize the flow field (Fig. 2b). Under these flow rate conditions, the ATP-containing Recording Medium of Pump A and the non-ATP-containing Recording Medium of Pump B were pulled into the branched cell introduction channel by the suction of Pump C, therefore, these media did not enter the cell culture chamber because the flow rates of Pumps A and B were very low. After culturing the cells for 17 hours, loading of Ca indicator to the cells was performed by replacing the reservoir pipette tip with a tip filled with Fluo 4-AM loading buffer, which was prepared by mixing Fluo 4-AM DMSO with Recording Medium at a concentration of 1 µg/µL. To introduce the loading buffer into the cell culture chamber, the flow rate of Pump C was increased to 1.0 µL/min in the same way as the cell culture (Fig. 2c). After 50 min to ensure loading and de-esterification of Fluo 4-AM, ATP stimulation was initiated by operating Pumps A and B (Fig. 2d). To prevent backflow from the cell introduction channel into the cell culture chamber, Pump C continued drawing at 0.05 µL/min, a flow rate low enough to be negligible for ATP stimulation.

The cells cultured in the microfluidic device were stimulated with ATP via two different modes to investigate the biological responses: pulse stimulation and stepwise stimulation. In the pulse mode, Pump A provided no flow and Pump B transported an ATP-free solution at the maximum flow rate (1.0 µL/min) initially. Then, Pump B was stopped and Pump A was started to deliver the ATP solution at the maximum flow rate (1.0 µL/min). After 120 s, Pump A was stopped and Pump B was started again at the maximum flow rate (1.0 µL/min). In this way, pulsed ATP flow was generated in the cell culture chamber. By contrast, a stepwise ATP flow waveform was generated by increasing or decreasing the flow rates of Pumps A and B at constant intervals. The flow rate of Pump A was increased by 0.25 µL/min from 0 to 1.0 µL/min at intervals of 120 s. The flow rate of Pump B was reduced by 0.25 µL/min from 1.0 to 0 µL/min at the same time intervals. Thus, the ATP concentration in the cell culture chamber rose along a stepwise waveform.

### Image processing

Quantitative image processing was used to extract single-cell responses from fluorescence live-cell imaging data. Based on the Otsu binarization method, we distinguished the cell region from the background region^31^. The average luminance value of the target cell area was taken as the cell response. The above image processing algorithm was performed using Cell Profiler software^32^.

### Hierarchical clustering

We performed hierarchical clustering of single-cell response data for ATP pulse stimulation (Fig. 7a). An M-by-N data matrix composed of N time points of M single-cell responses was normalized by scaling the variation to 1 in the row direction and averaging to 0 in both the row and column directions using the following equation:

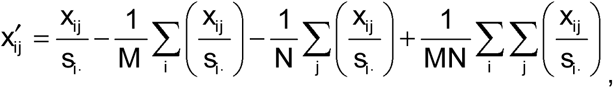

where si⋅, xij, and xi′ j represent the SD in the *i*th row of the raw matrix, the element of the raw matrix in the *i*th row and *j*th column, and the element of the normalized matrix in the *i*th row and *j*th column, respectively. We then implemented hierarchical clustering using the Euclidean distance for calculation of the intracluster distances and Ward’s method for calculation of the intercluster distances.

## Results and discussion

### Verification of mixing uniformity

The uniformity of mixing in the channel on the microfluidic device was verified by measuring the concentration of a fluorescent dye (FITC I dextran, 70 kDa) in the cell culture chamber. The CVs in the ROI of the cell culture chamber were measured for different flow rate ratios and are plotted in Figure 3. The CVs of the standard solutions, which were homogeneously premixed, were also measured and plotted on the same graph for each mixing ratio for reference. Over the entire range of mixing ratios, the measured CVs were <0.1, which is often used as the threshold for determining whether a solution is well mixed^29^. Likewise, the CVs of the solutions mixed in the mixing channel were very similar to the CVs of the standard solutions at all flow rate ratios.

**Fig. 3.**
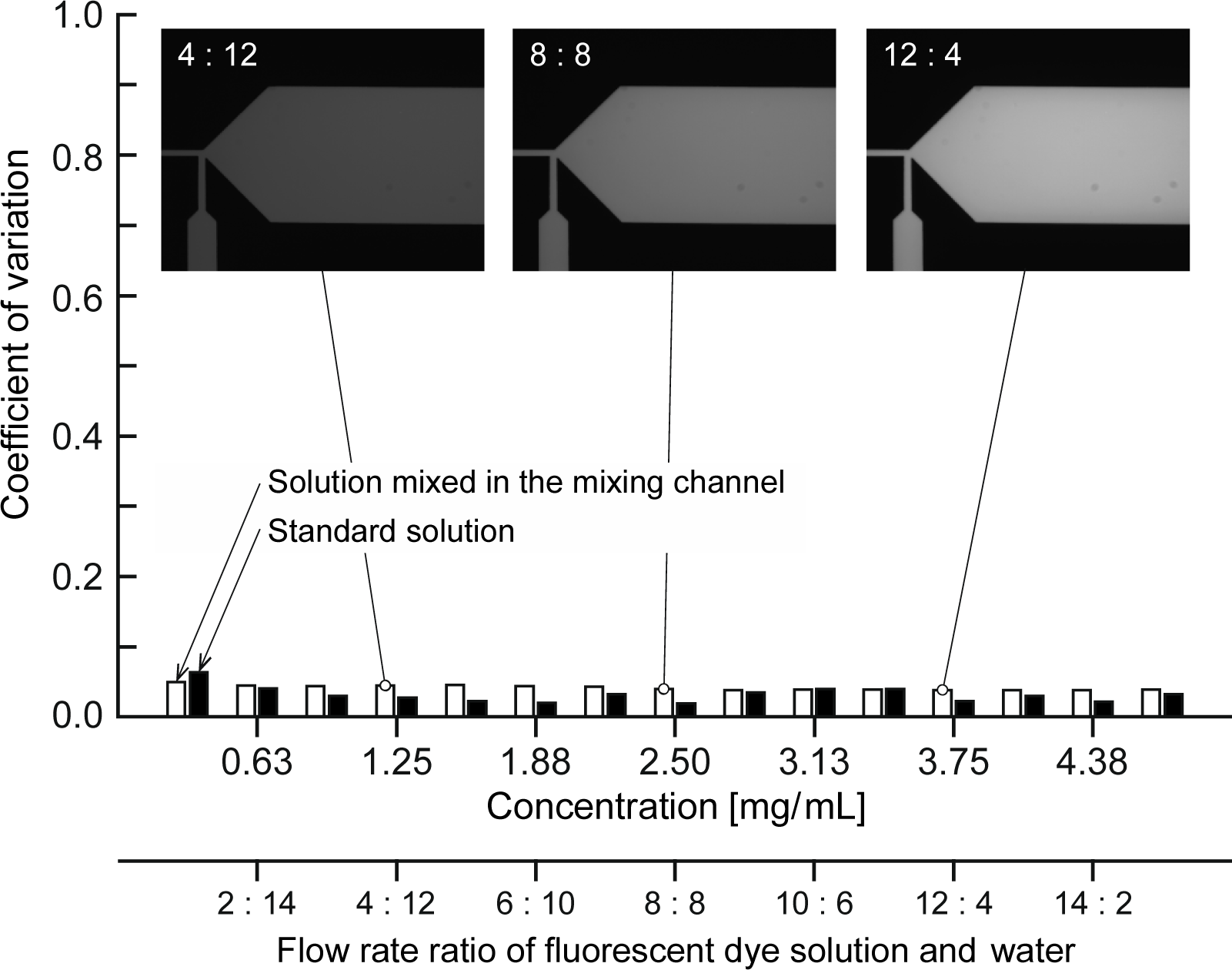
CV measurement using a fluorescence dye (FITC I dextran, 70 kDa). The CVs of the solutions mixed in the mixing channel (white bar) were sufficiently low and similar to those of the standard solutions (black bar), indicating that the solutions provided by Pumps A and B were mixed well at all mixing ratios.

A well-mixed solution should ideally result in a CV = 0 due to no variation in concentration across the ROI. However, the CVs of the standard solutions were not zero, because the current measurement system has a limited digital imaging resolution, resulting in noise in the captured images. Therefore, it is reasonable to conclude that the CVs of the solutions mixed in the mixing channel were sufficiently low at all mixing ratios and that the two solutions of Pumps A and B were mixed well in the cell culture chamber. The diffusion coefficient of the fluorescent dye (FITC I dextran, 70 kDa) used in this experiment is 23 µm^2^/s in water^33^. As a higher diffusion coefficient results in a shorter mixing time, the developed microfluidic device allows for molecules with a diffusion coefficient of ≥23 µm^2^/s to be distributed homogeneously in the cell culture chamber at the desired concentration.

### Temporal concentration regulation

Figure 4 shows a calibration curve of the relationship between dye (FITC I dextran, 70 kDa) concentration and fluorescence intensity. The measurement was repeated three times at each concentration. The calibration curve was nonlinear, and the gradient decreased gradually with increasing dye concentration. The measurement variation was greater at high concentrations compared with low concentrations.

**Fig. 4.**
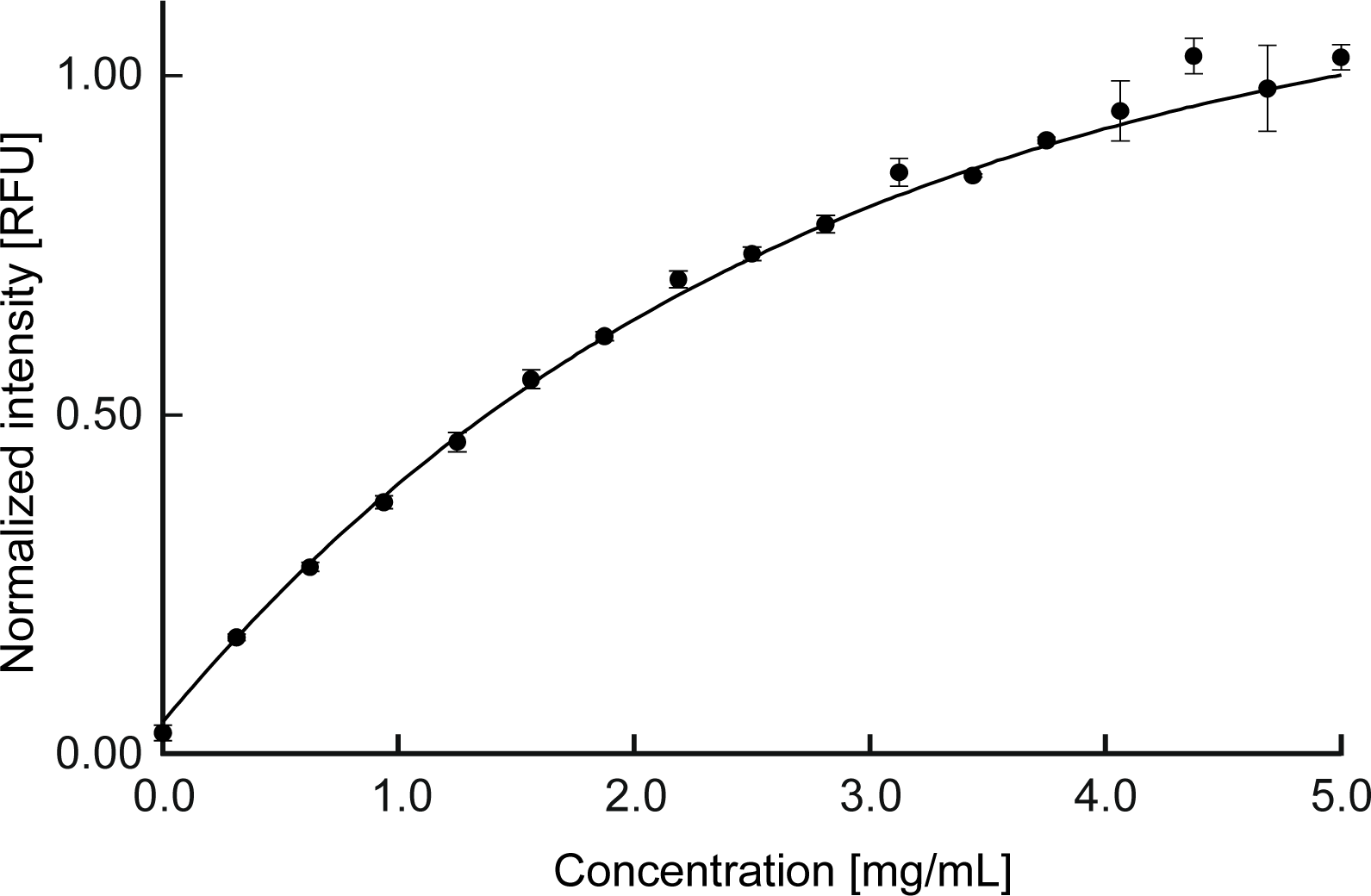
Relationship between dye (FITC I dextran, 70 kDa) concentration and fluorescence intensity. The fluorescence intensity was normalized. A calibration curve (solid line) was generated and plotted by fitting the measurement data (solid circles) to the exponential function.

The fluorescence intensity of a dye flowing in a microchannel usually shows a nonlinear increase with increasing dye concentration because of the thickness of the flow channel. The intensity of the excitation light decays due to interaction with other fluorescent molecules present in the light path before it reaches the fluorescent dye molecules. Meanwhile, the fluorescent light emitted by a fluorescent molecule can be blocked or absorbed by other fluorescent molecules present in the light path. Thus, in general, the fluorescence intensity detected by a CCD camera is not proportional to the concentration of dye. Variations in the three measurements conducted in this study could be attributed to unstable light intensity of the mercury lamp used as the excitation light source. It is difficult to maintain uniform light intensity of a mercury lamp over a long period of time. The relationship between the concentration and fluorescence intensity is expressed approximately by the following exponential function:

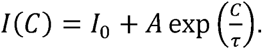

By fitting the measured data to this equation, the constants used in the equation were determined as follows: *I*_0_ = 1.1768, *A* = −1.1305, *τ* = −2.6854. The dye concentration can be derived from the measured fluorescence intensity using this equation and associated constants.

Two primary types of concentration waveforms (rectangular and triangular waves) were formed and measured in the cell culture chamber to demonstrate the performance of the developed system in terms of concentration regulation. Figure 5 shows the flow rate sequences of Pumps A and B and the results of time-series measurements of fluorescent dye concentration. Simple on-off switching of the dye solution and water flows at 90-s intervals allowed the dye concentration in the cell chamber to change periodically between the maximum and minimum values along a rectangular waveform (Fig. 5a). The time constant required for switching of the concentration was almost 40 s, which remained unchanged when the time interval was extended to 180 s (Fig. 5b). By gradually changing the flow rate ratio between the dye solution and water flows, the concentration in the cell chamber increased and decreased with a constant gradient along the triangular waveform (Fig. 5c). The gradient was kept constant when the time period was doubled (Fig. 5d). In both waveforms, the periodic change in concentration was very stable and highly reproducible for 12 min or longer. The concentration of dye in the cell chamber responded rapidly to changes in pump parameters and accurately reflected the programmed flow rate sequence. This indicates that the developed system has a remarkable capacity for forming various waveforms of concentration in the cell chamber without any complicated operation of the pumps.

**Fig. 5.**
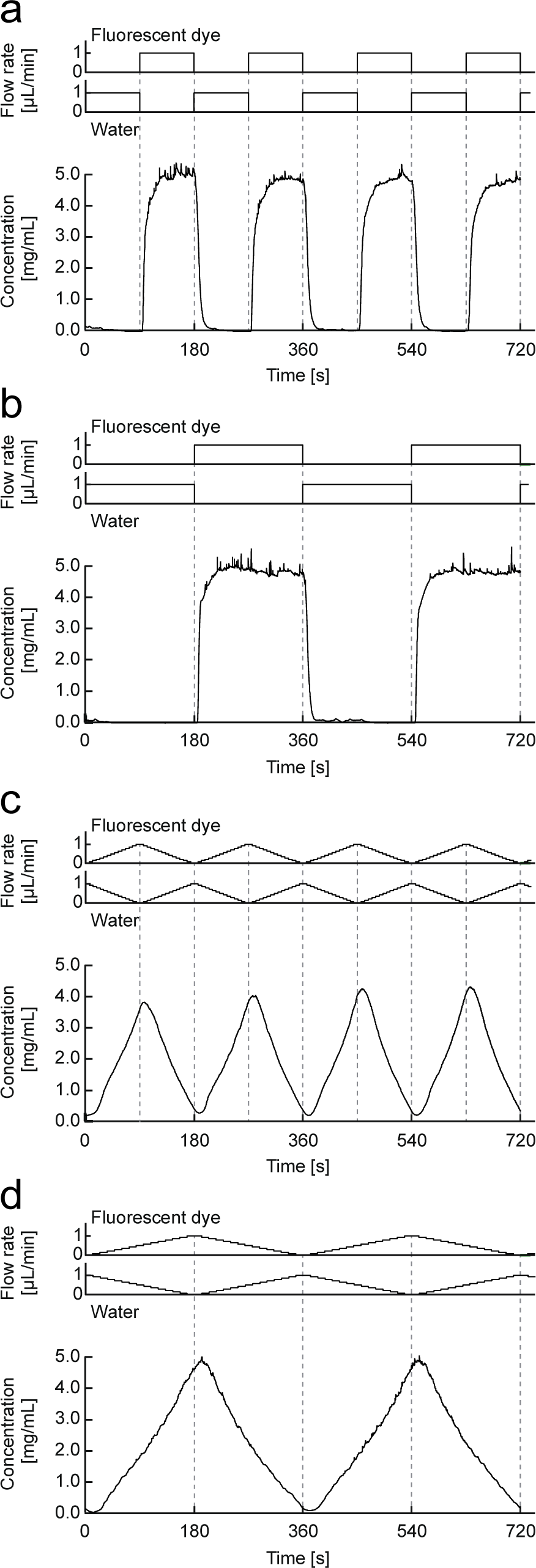
Flow rate sequences of Pumps A and B and time-series measurements of fluorescent dye concentrations. (a) On-off switching of dye solution flow and water flow at 90-s intervals and (b) 180-s intervals. (c) Repeating the upslope and downslope of the flow rates alternately at 90-s intervals and (d) at 180-s intervals.

### Cell stimulation and analysis

Changes in the Ca^2+^ concentration in CHO-K1 cells were measured when the cells were stimulated with ATP using the developed microfluidic cell culture system. Figure 6 shows the results of the cell simulation analyses. Bright-field microscopic images of CHO-K1 cells cultured in the device are shown in Figure 6a. The seeded cells immediately adhered to the collagen-coated surface of the cell culture chamber. After 17 h of culture, the cells proliferated and reached a confluent state. Figure 6b shows sequential fluorescence images of the cell culture chamber when the ATP solution was supplied to the cells in the pulse mode. The images in the upper part of the figure show red fluorescence (rhodamine B), which is representing the ATP concentration in the cell culture chamber. Although the green-colored excitation light was filtered out using a bandpass filter, some fluorescence of Fluo 4 emitted from the cells passed through and this fluorescent leak was recorded in the red fluorescence images. Green fluorescence (Fluo 4), which is representing the intracellular Ca^2+^ concentration, is shown in the lower part of the figure. The flow rate sequence, the ATP concentration in the cell culture chamber, and the fluorescence intensity of Fluo 4 are plotted in Figure 6c and d. When the cells were stimulated by ATP in pulse mode, the intracellular Ca^2+^ concentration spiked sharply in response to the rise in ATP concentration and then gradually decreased (Fig. 6c). Moreover, when the ATP concentration was increased in a stepwise manner, the cells showed several rapid increases intracellular Ca^2+^ concentration, in synchronization with the stepwise increases in ATP concentration (Fig. 6d). Additionally, the peaks of Ca^2+^ concentration spike was attenuated step by step.

**Fig. 6.**
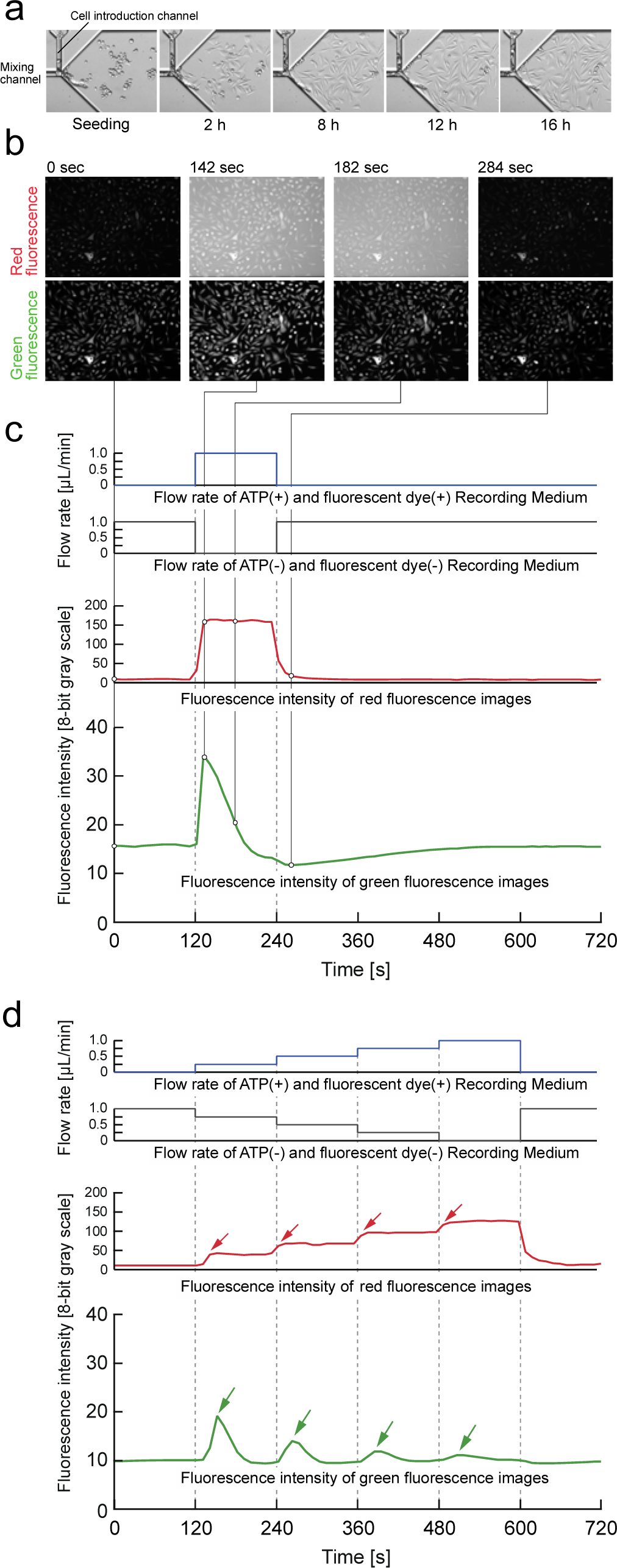
Cell stimulation with ATP and measurement of intracellular Ca^2+^ concentrations using CHO-K1 cells. (a) Microscopic images of CHO-K1 cells cultured in the device. (b) Sequential fluorescence images of the cell culture chamber when the ATP solution was applied to the cells in the pulse mode. The upper red fluorescence (rhodamine B) images show the ATP concentration. The lower green fluorescence (Fluo 4) images show the intracellular Ca^2+^ concentration. (c) The flow rate sequence, ATP concentration in the cell culture chamber, and fluorescence intensity of Fluo 4 when the cells were stimulated by ATP in pulse mode and (d) stepwise mode.

Although the ATP concentration around CHO-K1 cells was increased to the same level in both the pulse and stepwise stimulations, the cells exhibited different responses to each stimulation. This phenomenon suggests that intracellular Ca^2+^ concentration changes in response to temporal variations in ATP concentration rather than the concentration around cells. The observed attenuation of the peaks of intracellular Ca^2+^ concentration in the stepwise stimulation mode could be attributed to the exhaustion of Ca^2+^ ions stored in the endoplasmic reticulum.

### Hierarchical clustering of single-cell responses and quantitative analysis of response characteristics

The following quantitative analysis was performed to analyze the single-cell diversity of the Ca^2+^ response to ATP active stimulation by the microfluidic cell culture system described above. First of all, we implemented an image-processing method based on the Otsu binarization method to examine fluorescence imaging data and extract single-cell Ca^2+^ responses to pulse stimulation (see Experimental section). In order to quantify the diversity of time responses per cell, hierarchical clustering was performed on the temporal pattern of single-cell responses (Fig. 7a). The temporal patterns of Ca^2+^ response were classified into four clusters (Fig. 7b). We evaluated the maximum response value and fold change of each cell response per cluster in order to estimate the response characteristics to the stimulus (Fig. 7c). Clusters 1 and 2 showed the largest maximum response values, whereas Cluster 4 showed the smallest. By contrast, Cluster 4 showed the largest fold change, suggesting that Cluster 4 accurately represents the response despite exhibiting the lowest maximum response.

**Fig. 7.**
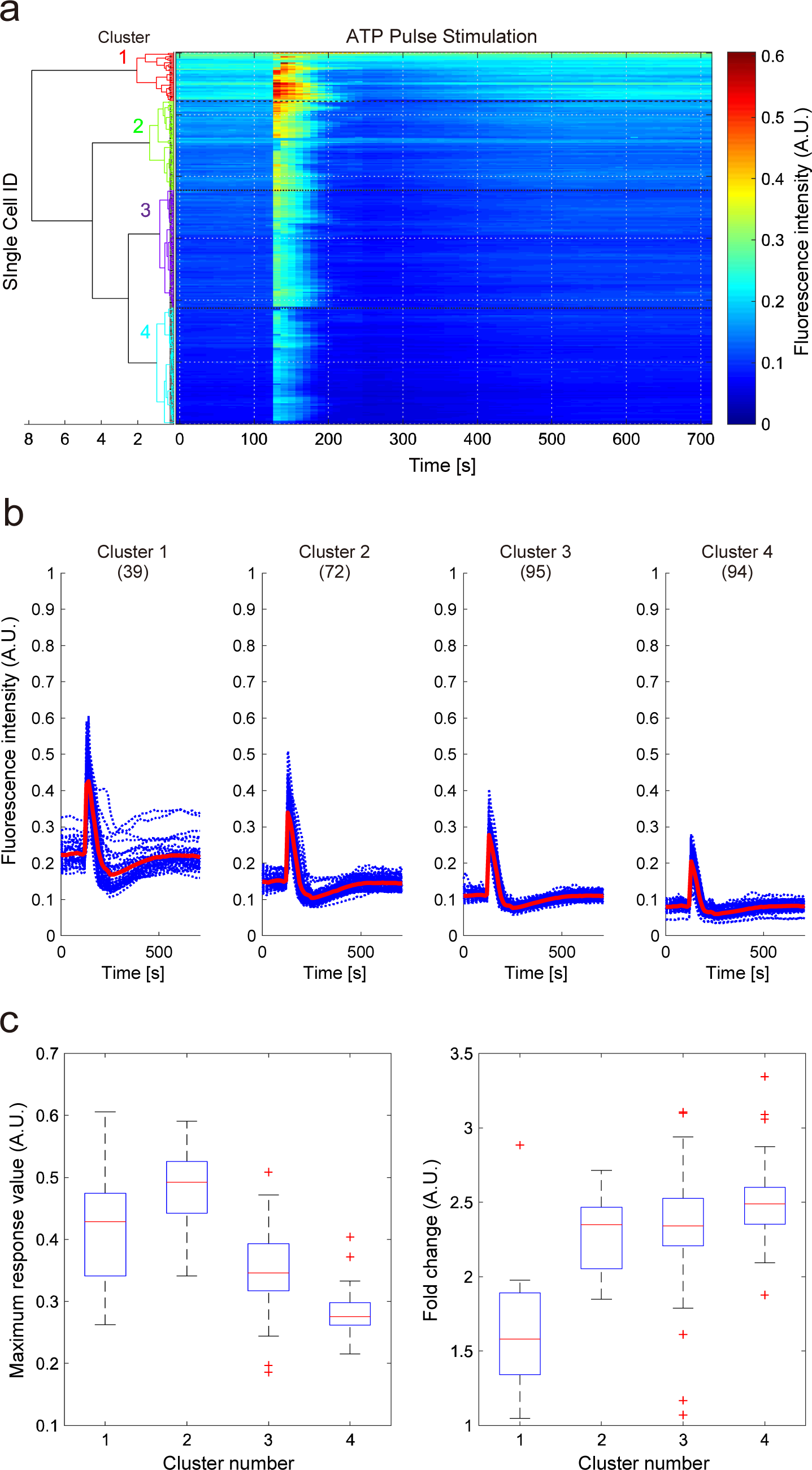
Hierarchical clustering of single-cell Ca^2+^ responses to ATP pulse stimulation and feature characteristics before and after stimulation of each cluster. (a) Dendrogram constructed by hierarchical clustering of single-cell Ca^2+^ responses to ATP pulse stimulation. Each row in the heatmap contains an individual cell response to ATP pulse stimulation. (b) Temporal patterns of cell responses to ATP pulse stimulation in each cluster. Blue and red lines indicate individual cell response and mean response, respectively. The number of cells is shown in parentheses. (c) Feature values for each cluster (upper: maximum response value; lower: fold change). Fold-change values were obtained by normalizing the maximum response value at a mean value of five points before stimulation.

The developed microfluidic system provides biological researchers with a new approach for dynamically and precisely controlling the biochemical environment around cells cultured in a small space, which would be impossible with traditional methods based on dish culturing and pipetting. The developed system enables us to apply a well-controlled biochemical stimulus to cells as the input and to acquire cell responses to the applied biochemical stimulus as the output. By measuring single-cell responses to actively controlled inputs, it is possible to perform clustering and modeling analyses accurately in order to quantify the diverse responses occurring at the single-cell level. From these results, it is confirmed that the developed system can be a powerful tool for use in investigating the biological responses of cells, including intracellular signal transduction.

## Conclusions

A microfluidic cell culture system was developed for use in dynamic cell signaling studies. The developed system enables varying patterns of signal molecule concentrations over time using pulse and triangular waveforms for stimulating cells in a time-dependent manner. Cells cultured in the device and their responses to stimuli are visualized and measured using time-lapse fluorescence microscopy imaging.

The performance of the developed system in concentration regulation was demonstrated and evaluated by measuring the uniformity of mixing and the formation of two primary types of concentration waveforms, rectangular and triangular. The CVs of measurement of the solutions were sufficiently low at all mixing ratios. This result demonstrated that the mixing channel of the developed microfluidic device, which has a high aspect ratio, enhances mixing and allows molecules with a diffusion coefficient of ≥23 µm^2^/s to be distributed homogeneously at a desired concentration in the cell culture chamber. By regulating the flow rate ratio between two syringe pumps in a programmed sequence, the concentration of a fluorescent dye could successfully be regulated along rectangular and triangular waveforms. The concentration in the cell chamber changed rapidly in response to the changes in pump operation parameters and accurately reflected the programmed flow rate sequence.

The developed system was applied to the measurement of intracellular Ca^2+^concentrations in CHO-K1 cells using a green fluorescent Ca^2+^ probe (Fluo 4). The cells showed rapid increases of intracellular Ca^2+^ level in synchronization with ATP stimulation. Furthermore, we implemented quantitative image analysis of fluorescence data and extracted single-cell Ca^2+^ responses to ATP pulse stimuli. Hierarchical clustering and quantitative analysis of single-cell data were performed and quantified the diversity of Ca^2+^ responses to ATP stimulation.

These results demonstrated that the developed microfluidic cell culture system is useful for the study of cellular signaling mechanisms and will contribute to investigations on the cellular responses such as intracellular signaling and to other time-based biological analyses of cells. Furthermore, cellular response data for active stimulation is essential for quantitative biology and data driven system biology research^34,35^. The developed microfluidic device has a potential to greatly contribute to these fields of research.

## Acknowledgements

This work was supported by CREST (JPMJCR12W3), JST, and partially by JSPS KAKENHI Grant Number JP26650075.

